# Ectomesenchymal identity emerges via relief of *Twist1* transcript destabilization

**DOI:** 10.1101/2025.09.02.673848

**Authors:** Lara C. Busby, Jessica R. Patrick, Luke W. Lyons, Megan L. Martik

## Abstract

During vertebrate development, cranial neural crest cells (CNCCs) differentiate into a variety of derivatives, including ectodermal cell types (neurons, glia, and pigment cells) as well as a suite of derivatives that are classically associated with the mesoderm (cartilage, bone and muscle) and are collectively termed ‘ectomesenchyme’. While the molecular decisions that guide CNCCs toward ectomesenchymal identity remain incompletely understood, the transcription factor Twist1 plays a central role. Here, we investigate the regulation of *Twist1* expression in CNCCs and find that *Twist1* is expressed by late migratory ectomesenchymal CNCCs in *Gallus gallus* and *Danio rerio* embryos. Using Hi-ChIP, ATAC-seq, and CUT&RUN sequencing data, we identify a distal enhancer for *Twist1* within the *Hdac9* locus that is active in the neural tube and CNCCs. Notably, this enhancer is directly bound by TFAP2 transcription factors and is active in pre-migratory CNCCs, a stage when *Twist1* transcripts are not detectable in CNCCs. We reconcile this temporal discrepancy by showing that the *Twist1* 3’ UTR of multiple vertebrate species (but not the non-vertebrate chordate *Ciona intestinalis*) is sufficient to destabilize GFP transcripts in the neural tube and surface ectoderm. Together, these findings reveal a vertebrate-specific, two-tiered regulatory mechanism that uncouples enhancer activity from transcript accumulation, gating the onset of *Twist1* expression in CNCCs and the acquisition of ectomesenchymal identity in vertebrate CNCCs.

## Introduction

The evolution of the vertebrate phylum is intimately associated with the emergence of a novel embryonic cell type, the neural crest (NC), which gives rise to a remarkable variety of cell type derivatives during embryogenesis (Martik and Bronner, 2021). Despite their ectodermal origin in the neural plate border, neural crest cells (NCCs) give rise to both ectodermal derivatives as well as cell types historically associated with the mesoderm, including chondrocytes, smooth muscle cells, and cardiomyocytes. These NCCs that form mesoderm-like derivatives are termed ectomesenchyme, reflecting their origin in the ectoderm yet acquisition of mesenchymal characteristics (De Beer, 1947; Horstadius, 1950; Platt, 1893). Though initially controversial, the contribution of NCCs to mesenchymal derivatives is now widely accepted and recognized as a key evolutionary innovation that enabled the expansion of the vertebrate head and adoption of a predatory lifestyle (Gans and Northcutt, 1983).

Seminal fate-mapping studies in a variety of vertebrate models have shown that the ability to give rise to some derivatives is restricted to subpopulations of NCCs that arise at distinct positions along the anteroposterior (AP) axis, a phenomenon termed axial regionalization. In amniotes, only cranial neural crest cells (CNCCs), which arise anterior to rhombomere 6, give rise to chondrocytes and osteocytes that form the majority of the craniofacial skeleton, including the jaw. Classical grafting experiments demonstrated that trunk NCCs transplanted to the cranial region do not contribute to cartilage of the developing head (Chibon, 1967; Lwigale et al., 2004; Nakamura and Ayer-le Lievre, 1982). However, CNCCs transplanted into the trunk region can form ectopic cartilage nodules (Le Douarin and Teillet, 1974). These findings suggest that there is an intrinsic difference in the potency of NCCs along the AP axis that is established prior to their emigration from the neural tube.

More recent work has demonstrated that these differences in potency are controlled by the local activation of gene regulatory subcircuits along the AP axis. Genes encoding a suite of cranial-specific transcription factors (*Brn3c, Lhx5, Dmbx1, Sox8, Tfap2b* and *Ets1*) are expressed by premigratory CNCCs, but not by premigratory trunk neural crest cells in amniotes (Simoes-Costa and Bronner, 2016). When three of these cranial transcription factors (*Sox8, Tfap2b*, and *Ets1*) were overexpressed in trunk NCCs, reprogrammed trunk NCCs were able to give rise to chondrocytes upon transplantation to the head environment, suggesting that these early cranial identity genes are sufficient to imbue trunk NCCs with ectomesenchymal potential (Simoes-Costa and Bronner, 2016). Subsequently, this cranial-specific NC GRN was shown to have arisen progressively throughout vertebrate evolution with gradual addition of network components in gnathostome lineages (Martik et al., 2019).

Arising in the dorsal neural tube, ectomesenchymally fated CNCCs migrate ventrally beneath the surface ectoderm into the pharyngeal arches, where late in their migration they undergo a transcriptional switch. This switch involves downregulating the expression of early migratory NC genes (such as *Sox10* and *Foxd3*) while upregulating the expression of ectomesenchymal genes (including *Dlx2*) (Blentic et al., 2008), and is dependent on arch-derived extrinsic signals, including FGF. Inhibition of FGF signaling disrupts this switch, causing CNCCs to inappropriately retain expression of *Sox10* and *Foxd3* in the pharyngeal arches (Blentic et al., 2008). Thus, previous work has hinted at key features of the intrinsic CNCC transcriptional state at premigratory stages, as well as aspects of the extrinsic signals from the head environment that are important for the emergence of ectomesenchyme. However, we currently lack a full understanding of the molecular mechanisms by which cells adopt ectomesenchymal identity.

In this study, we dissect the regulatory mechanisms that govern the acquisition of ectomesenchymal identity in CNCCs by focusing on the regulation of *Twist1. Twist1* has been implicated as a key node in the ectomesenchymal CNCC gene regulatory network (GRN) from single-cell RNA sequencing of sorted mouse CNCCs, where it was found to drive the decision between neuronal and ectomesenchymal cell fate (Soldatov et al., 2019). Overexpression of *Twist1* in trunk NC is sufficient to redirect the migratory path of these cells away from the dorsal root ganglion (toward the dermis) and activates the expression of *Prrx2*, another key mesenchymal identity gene (Soldatov et al., 2019). Additionally, numerous experiments in multiple vertebrate species have shown that loss of *Twist1* function in CNCCs results in a failure to activate ectomesenchymal gene programs and leads to maintained expression of early NC genes, in some cases biasing cells toward neuronal identity (Soo et al., 2002; Vincentz et al., 2007; Das and Crump, 2012; Kim et al., 2024). Notably, this phenocopies the consequences of loss of FGF signaling from the pharyngeal arches (Blentic et al., 2008), suggesting that *Twist1* may act upstream of other ectomesenchymal identity genes.

Here, we show that in avian and zebrafish embryos, ectomesenchymal identity emerges late during the migration of CNCCs to their destinations in the ventral head. Using previously published Hi-ChIP, CUT&RUN, and ATAC-seq data from avian CNCCs, we identify a distal enhancer that contacts the *Twist1* locus and drives robust reporter gene expression in the neural tube and migratory CNCCs. Surprisingly, this enhancer is active well before endogenous accumulation of the *Twist1* transcript, suggesting that there is an additional level of regulation of *Twist1* transcripts. We resolve this paradox by showing that the *Twist1* 3’ UTR potently destabilizes GFP transcripts in the neural tube and early migratory CNCCs, indicating that post-transcriptional regulation is a key contributor to the emergence of ectomesenchyme.

Together, our findings reveal a previously unrecognized, two-tiered regulatory mechanism, coupling enhancer priming and transcript destabilization, that ensures *Twist1* activation occurs only once CNCCs reach the ventral head environment. This mechanism not only explains how ectomesenchymal identity is precisely controlled in space and time but also illuminates a potential route by which the neural crest GRN evolved during vertebrate evolution to give rise to ectomesenchyme.

## Results

### *Twist1* is expressed by late migratory ectomesenchymal neural crest cells in the vertebrate embryo

To characterize the emergence of ectomesenchymal cell states in space and time, we performed multiplexed in situ hybridization (Choi et al., 2018) to detect transcripts of the ectomesenchyme-associated gene *Twist1*, as well as NC specifier genes *Sox10* and *Ets1*. Co-staining for NC specifier gene expression was crucial for interpretation in these analyses because *Twist1* is broadly expressed in mesodermal tissues across vertebrates during development.

In chicken embryos, *Twist1* expression is detected in the cranial mesoderm from premigratory CNC stages (HH8), but not in premigratory CNCCs themselves, which are marked by *Sox10* and *Ets1* expression (*Figure 1A, Figure S1*). By HH10, however, late migratory CNCCs close to their destinations begin to upregulate expression of *Twist1* (*Figure 1B-C*). In the anterior mesenchyme overlying the forebrain, late migratory CNCCs in close apposition to the optic vesicle express *Twist1* (*Figure 1B*). More posteriorly, *Sox10*/*Ets1*-positive CNCCs migrate ventrally beneath the surface ectoderm (*Figure 1C*), with *Twist1* transcripts only becoming detectable in the most ventral cells, furthest along their migratory path (arrowheads in *Figure 1C*). Thus, in chicken embryos, the key ectomesenchymal identity gene *Twist1* is not activated in CNCCs until they are late in migration.

**Figure 1:**
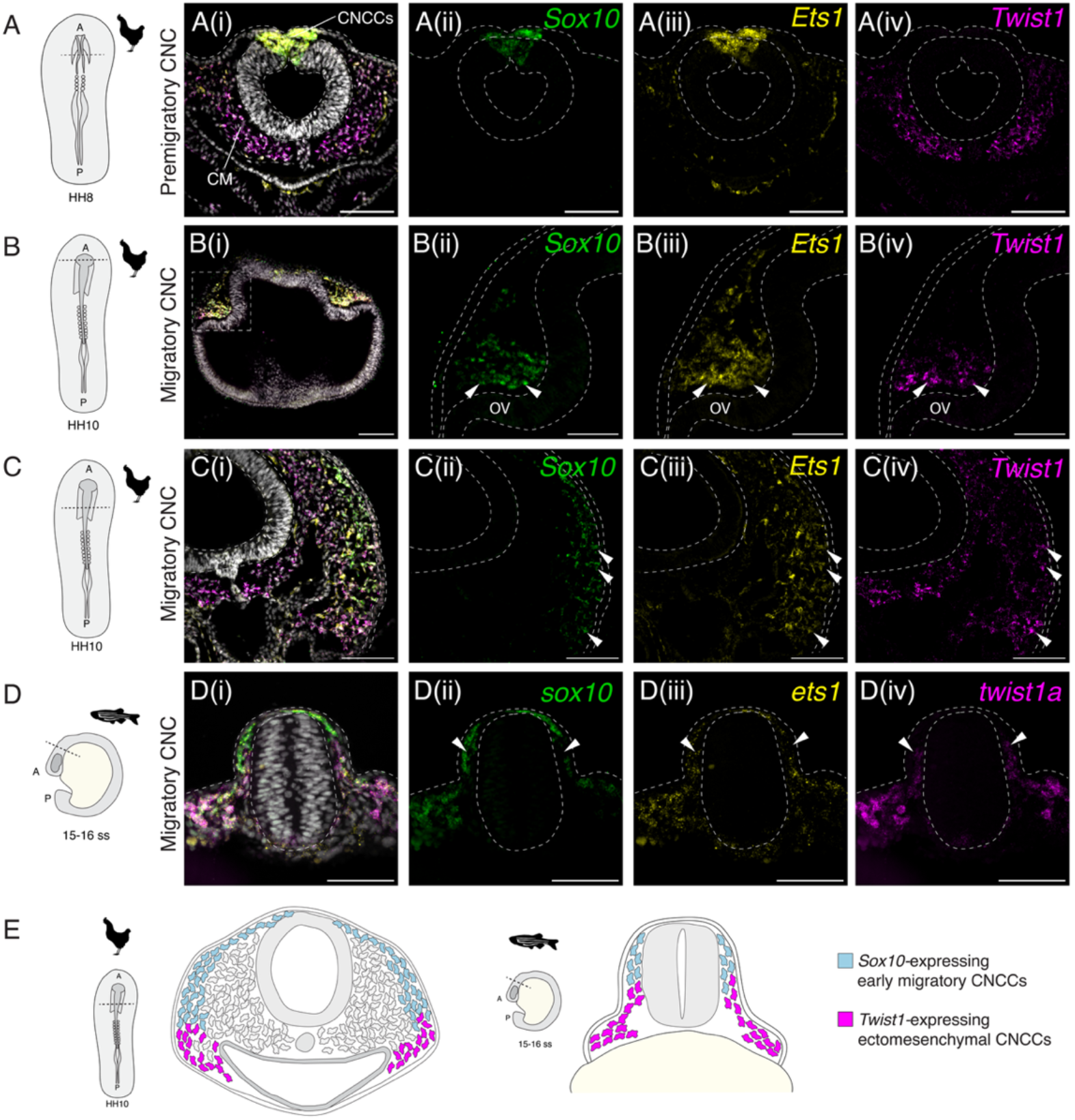
Twist1 is expressed by late migratory ectomesenchyme. (A)-(D) are summed slice intensity projections of multiplexed HCR staining for transcripts of the early neural crest specifier gene *Sox10*, the cranial NC transcription factor *Ets1* and the ectomesenchymal identity gene *Twist1*. In each panel, (i) shows a composite image with DAPI, *Sox10, Ets1* and *Twist1* transcripts, and (ii)-(iv) are single channel images for *Sox10, Ets1* and *Twist1* respectively. (A)-(C) Staining in chicken embryo transverse sections. (A) shows that premigratory CNCCs in the dorsal neural tube at HH8 express both *Sox10* and *Ets1* but do not express *Twist1*, whilst the underlying cranial mesoderm (CM) expresses *Twist1* at this stage. (B)-(C) Sections in chicken embryos at HH10 show that *Twist1* is only upregulated in late migratory CNCCs (B) in the forebrain region overlying the prospective optic vesicle (OV) and (C) more posteriorly within the cranial region in cells migrating sub-ectodermally into the pharyngeal arches. (D) Zebrafish embryo cranial region section at 15-16ss stained for *sox10, ets1* and *twist1a*. (E) Schematic summarizing *Twist1* expression analyses in chicken and zebrafish. *Animal silhouettes are from Phylopic, scale bars in (A), (B(i)) and (C) represent 100 µm, and in (B(ii)-(iv)) and (D) represent 50 µm*.

To test whether this spatiotemporal dynamic is conserved, we stained *Danio rerio* embryos for *sox10, ets1* and *twist1a* transcripts at stages between 4 somite stage (4ss) and 24 hours post fertilization (hpf). Consistent with previous reports (Das and Crump, 2012; Germanguz et al., 2007; Tatarakis et al., 2021), *twist1a* transcripts become detectable in CNCCs as they migrate over the developing eye, as well as in late migratory and post-migratory CNCCs located in the pharyngeal arches (*Figure 1D, Figure S2*).

In summary, we find that across the vertebrate phylum, representative species of multiple lineages express *Twist1* in late migratory ectomesenchyme of the head (*Figure 1E*). This expression challenges the long-standing view of *Twist1* as a driver of epithelial-to-mesenchymal transition in neural crest cells and instead establishes *Twist1* as a key player in the ectomesenchymal cell fate decision and a regulatory node worth dissecting further.

### Identification of a *Twist1* enhancer in the *Hdac9* locus that drives reporter gene expression in the neural tube and migratory CNCCs

To ask how *Twist1* gene expression is initiated in ectomesenchymal NC, we combined sequence conservation analysis with existing DNA-DNA contact (H3K27ac Hi-ChIP), chromatin accessibility (ATAC-seq) and Cut & Run (TFAP2A, TFAP2B, H3K27ac) datasets from sorted avian CNCCs to identify putative enhancer sequences (Hovland et al., 2020; Azambuja & Simoes-Costa, 2021). We find that the *Twist1* promoter region forms numerous distal genomic contacts in dorsal neural fold cells, many within the neighboring *Hdac9* locus (*Figure 2A, Supplementary Table 1*), though none of these loops are differentially enriched in CNCCs relative to whole embryo samples. Altogether, we used enhancer reporter assays to test a total of eleven putative enhancer sequences (*Supplementary Table 2*) and found two sequences with mesodermal activity (*Figure S3*) and one sequence that drove activity in the neural crest (*Figure 2C-F*).

**Figure 2:**
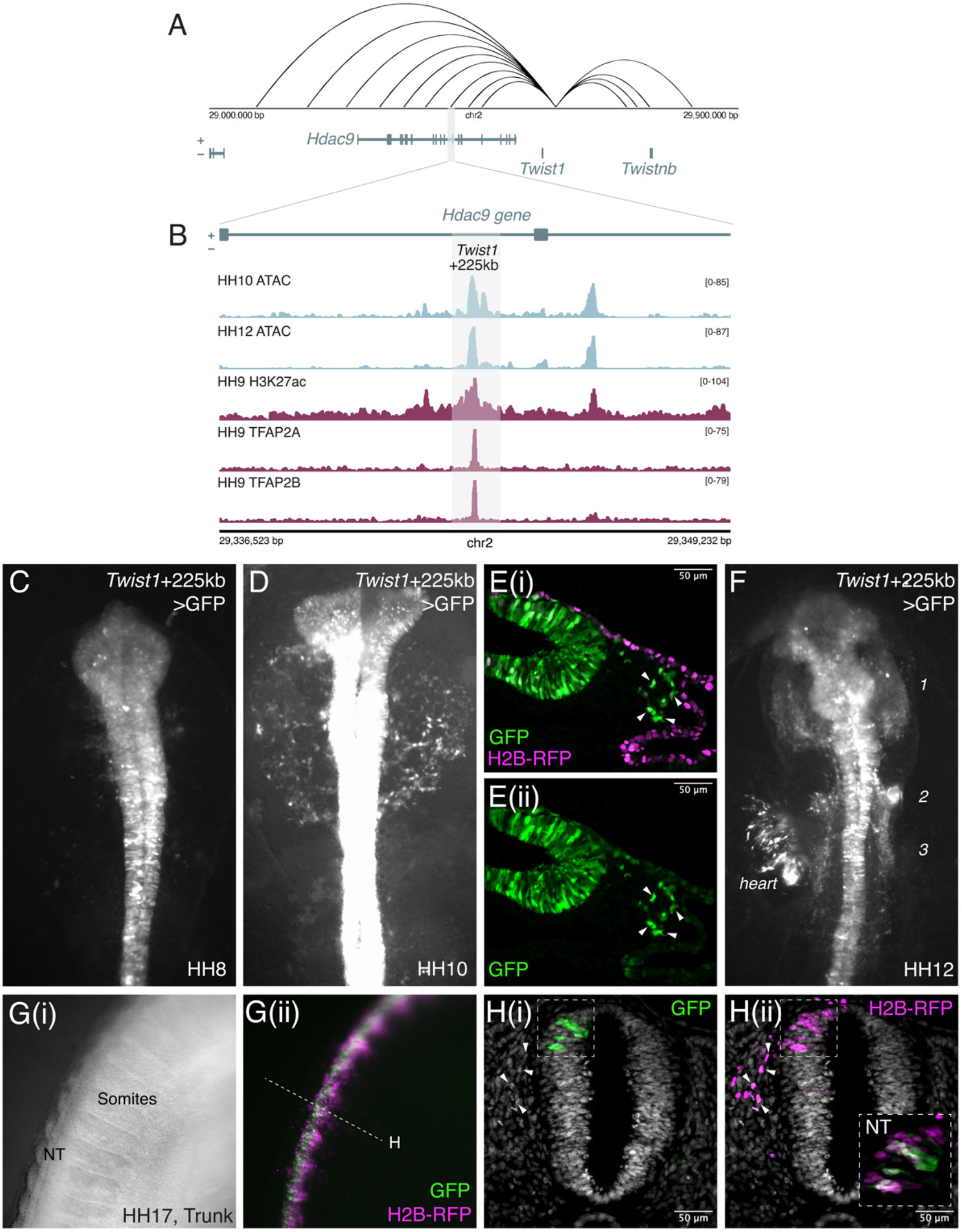
Identification of an enhancer element driving Twist1 expression in cranial neural crest cells. (A) Contact map showing distal genomic contacts of the *Twist1* gene promoter region; the region containing *Twist1*+225kb is highlighted by the grey box. Raw data derived from Azambuja et al., 2021. (B) Multi-modal tracks showing chromatin accessibility (Hovland et al., 2022), H3K27 acetylation (Bhattacharya et al., 2020) and TFAP2A/2B genome occupancy (Rothstein and Simoes-Costa, 2020) in the region of the *Twist1*+225kb enhancer, located within an intron of the *HDAC9* gene. (C), (D) & (F) GFP fluorescence driven by the *Twist1*+225kb: GFP enhancer reporter at HH8, HH10 and HH12 respectively. Numbers in (F) represent the three streams of cranial neural crest. (E(i) and (ii)) Representative transverse section showing *Twist1*+225kb: GFP fluorescence and H2B-RFP (electroporation control) at HH10. Arrowheads indicate migrating CNCCs with enhancer activity. (G)-(H) *In ovo* electroporations of the *Twist1*+225kb: GFP enhancer reporter injected into the neural tube at HH10-12. (G(i)) Shows a brightfield image of the embryonic trunk at HH17 and (G(ii)) shows the corresponding composite image for GFP and H2B-RFP fluorescence. (H) Trunk section of *in ovo* electroporated embryo shows that GFP expression is driven in the neural tube by *Twist1*+225kb but despite labelling of trunk NCCs emigrating from the tube (arrowheads in (ii)) no GFP activity is present in these cells, suggesting that activity of the *Twist1*+225kb enhancer is cranial-specific in NCCs.

We identified a 709bp sequence within intron 13 of the *Histone Deacetylase 9* (*Hdac9*) gene that drives robust reporter expression in the neural tube and in migrating cranial and vagal NC, herein termed *Twist1*+225kb (*Figure 2B-F*). This genomic sequence is accessible, acetylated, and directly bound by the transcription factors TFAP2A and TFAP2B in premigratory CNCCs, supporting its role as an enhancer *(Figure 2B*). Using an enhancer reporter assay, we found that *Twist1*+225kb drives expression of GFP throughout the neural tube and in emigrating CNCCs at HH8-HH10 (*Figure 2C-E*). By HH12, *Twist1*+225kb labels migratory CNCCs in the three primary CNCC streams (*Figure 2F*) (Theveneau and Mayor, 2012).

Given the cranial restriction of *Twist1* expression in neural crest cells, we asked whether *Twist1*+225kb activity was similarly restricted. Interestingly, we find that *Twist1*+225kb drives reporter expression in trunk neural tube cells but not in migrating TNCCs (*Figure 2G-H*). Therefore, the activity of this enhancer recapitulates the cranial restriction of *Twist1* expression and ectomesenchyme emergence to the cranial region.

Taken together, our results thus far uncover a distal enhancer that contacts the *Twist1* promoter and is active in premigratory CNCCs, well before *Twist1* transcripts are detectable (*Figure 1*). This striking discrepancy between enhancer activity and mRNA expression suggests an additional regulatory layer to precisely control *Twist1* presence and emergence of ectomesenchymal identity. We therefore asked what upstream factors could be controlling the activity of this enhancer and whether their activity could account for the disconnect between enhancer activation and *Twist1* transcript abundance.

### The *Twist1*+225kb enhancer is regulated by canonical neural crest transcription factors

To identify upstream transcription factors that regulate the *Twist1*+225kb enhancer, we used FIMO (Grant et al., 2011) to predict transcription factor binding sites, which revealed the presence of putative binding sites for TFAP2, FOXD, TGIF1, MAF, ZIC, PRDM, and POUV/OCT4 (*Figure 3A*). Each of these transcription factors has previously been implicated in neural crest development. FOXD3 and TFAP2A are pan-NC transcription factors that are expressed in premigratory and migratory NCCs at all axial levels, and TFAP2B is a cranial-specific transcription factor (Simoes-Costa & Bronner, 2016; Rothstein & Simoes-Costa, 2020). TFAP2A and TFAP2B have been shown to heterodimerize in CNCCs and act as pioneer transcription factors (Rothstein & Simoes-Costa, 2020). TGIF1, MAFA, and MAFB have been associated with the cardiac NC (Dhillon-Richardson et al., 2025; Gandhi et al., 2020a). HMGA1 is a chromatin remodeler active in the ectoderm including CNCCs (Gandhi et al., 2020b). POUV/OCT4 has been implicated in the ectomesenchymal potential of CNCCs and is maintained or reactivated in the ectoderm (Buitrago-Delgado et al., 2015; Zalc et al., 2021; Pajanoja et al., 2023).

**Figure 3:**
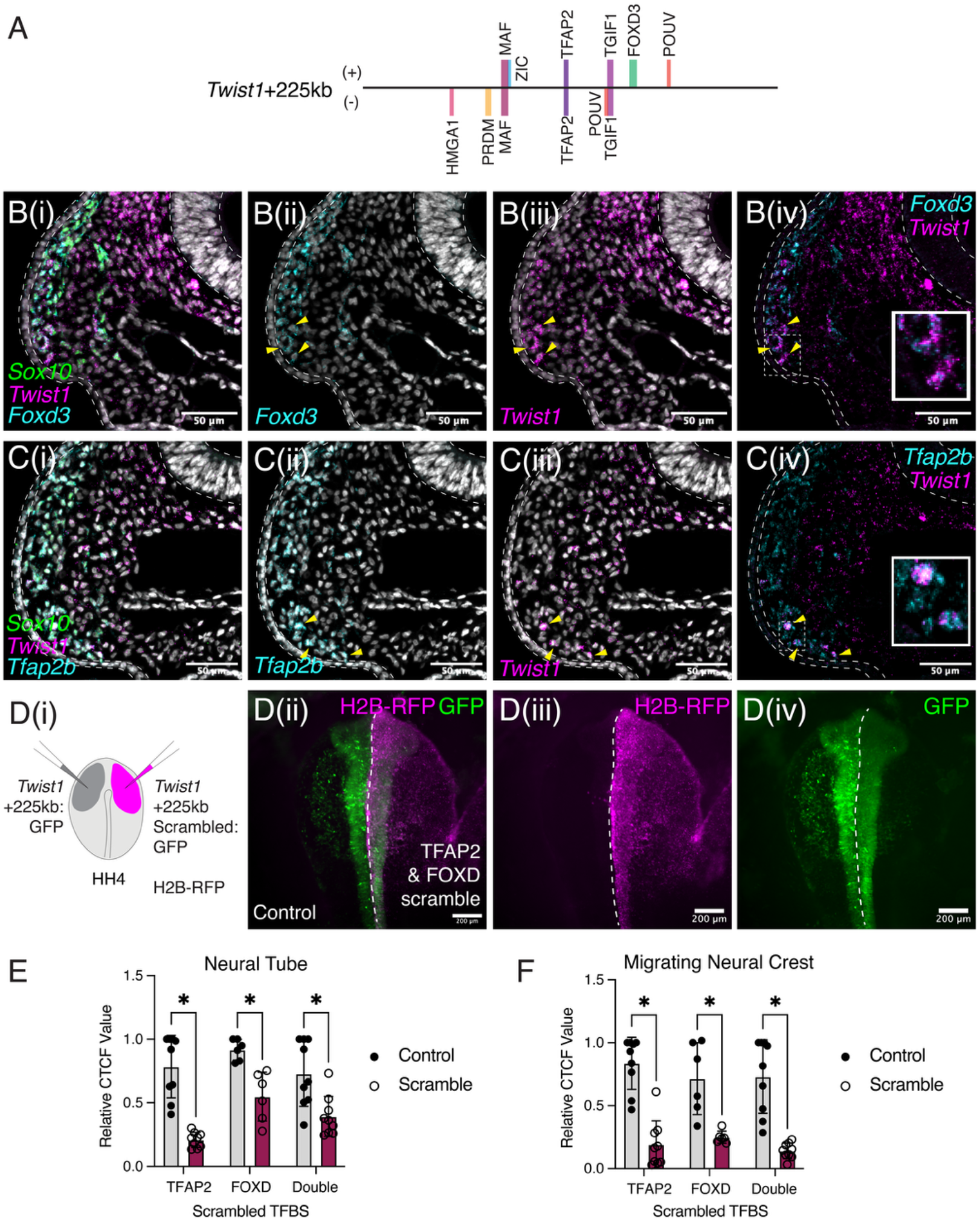
The Twist1+225kb enhancer is regulated by canonical neural crest transcription factors. (A) Box diagram of the *Twist1*+225kb enhancer sequence with key predicted transcription factor binding sites annotated to scale. (B) Multiplexed HCR staining for *Foxd3* and *Twist1* at HH10 shows co-expression of these two transcripts in late migratory CNCCs. Arrowheads indicate cells co-expressing *Foxd3* and *Twist1*. (C) Multiplexed HCR staining for *Tfap2b* and *Twist1* at HH10 shows co-expression of these two transcripts in late migratory CNCCs. Arrowheads indicate cells co-expressing *Tfap2b* and *Twist1*. (D) (i) Schematic of scrambled enhancer experiment. The *Twist1*+225kb: GFP scrambled construct shown in (D) has both the TFAP2 and FOXD binding sites scrambled. (ii) Composite GFP and RFP fluorescence image of representative injected embryo at HH10. (iii) H2B-RFP fluorescence - electroporation control showing cells which received the *Twist1*+225kb: GFP scrambled construct. (iv) GFP fluorescence. (E) – (F) Quantification of GFP fluorescence intensity (corrected total cell fluorescence, CTCF) in regions of the neural tube (E) or lateral to the neural tube (F) reveal significantly reduced fluorescence resulting from scrambling of TFAP2, FOXD or TFAP2 & FOXD motifs in the *Twist1*+225kb enhancer sequence, suggesting that these motifs are important for enhancer activity. *Statistics calculated using unpaired t-tests, with the following p-values: neural tube TFAP2 scramble 0*.*000004, neural tube FOXD scramble 0*.*001505, neural tube double scramble 0*.*004071, migrating NC TFAP2 scramble 0*.*000003, migrating NC FOXD scramble 0*.*002773, and migrating NC double scramble 0*.*000021*.

To test whether the spatiotemporal expression of these putative regulators is consistent with their regulation of *Twist1*, we performed HCR against *Sox10, Twist1* and either *Tfap2b* or *Foxd3*. We observed the coexpression of both *Tfap2b* and *Foxd3* transcripts with *Twist1* in ventral late migratory CNCCs at HH10 (*Figure 3B-C*), suggesting that these transcription factors could have a functional role in driving *Twist1* expression in CNCCs. Notably, *Tfap2b* and *Foxd3* transcription factors begin to be expressed when neural crest cells are premigratory in the dorsal neural tube, much earlier than *Twist1* transcripts become detectable, suggesting they could prime *Twist1* activation via the *Twist1*+225kb enhancer before transcripts accumulate.

To directly test the functional importance of a subset of these binding sites in driving *Twist1*+225kb enhancer activity, we generated modified enhancer constructs, each with the predicted binding motif for a transcription factor scrambled. To allow comparison of relative enhancer activity after scrambling binding sites, we contralaterally electroporated the control enhancer construct and the scrambled construct with a H2B-RFP-encoding plasmid (*Figure 3D(i)*) and then assessed GFP fluorescence at HH10 (*Figure 3D*). We found that constructs with a single transcription factor binding site scrambled, either TFAP2 or FOXD, led to significantly reduced activity of the enhancer in migratory CNCCs and the neural tube (*Figure 3E, F*). Strikingly, scrambling both sites abolished enhancer activity in the migrating CNCCs relative to the control construct (*Figure 3D(iv), F*). Together, these results suggest that the binding of TFAP2 and FOXD transcription factors to the *Twist1*+225kb sequence is essential for enhancer activity in neural crest cells.

Despite the robust enhancer activation via canonical neural crest transcription factor binding, *Twist1* transcripts remain undetectable until much later in migration. This paradox, therefore, challenges the conventional assumption that enhancer activity and transcript accumulation are tightly coupled. To resolve this, we considered that additional layers of regulation may gate the precise timing of *Twist1* expression and therefore the emergence of ectomesenchymal identity.

### The *Twist1* transcript is post-transcriptionally destabilized in the ectoderm

Given the clear difference in timing between onset of *Twist1*+225kb enhancer activity and the accumulation of *Twist1* transcripts (late migratory CNCCs), we asked whether *Twist1* transcripts may be subject to post-transcriptional regulation. Post-transcriptional regulation of a transcript commonly occurs via sequence and/or structural elements found in the 3’ untranslated region (3’ UTR). To test the influence of the *Twist1* 3’ UTR on transcript stability, we cloned a construct consisting of a ubiquitous promoter driving the expression of destabilized GFP followed by the chicken *Twist1* 3’ UTR (*Figure 4A*). As a control, we electroporated an equivalent construct with the *Beta-globin* 3’ UTR (*Figure 4A*), which is known to stabilize transcripts in a variety of cellular contexts (Hutchins et al., 2020; Russell and Liebharber, 1996). To enable direct side-by-side comparison of GFP fluorescence as a readout of transcript stability, we electroporated the *Twist1* 3’ UTR construct (with a H2B-RFP plasmid as an electroporation control) on the right of an embryo and the *Beta-globin* 3’ UTR construct on the left side (*Figure 4A-B*).

**Figure 4:**
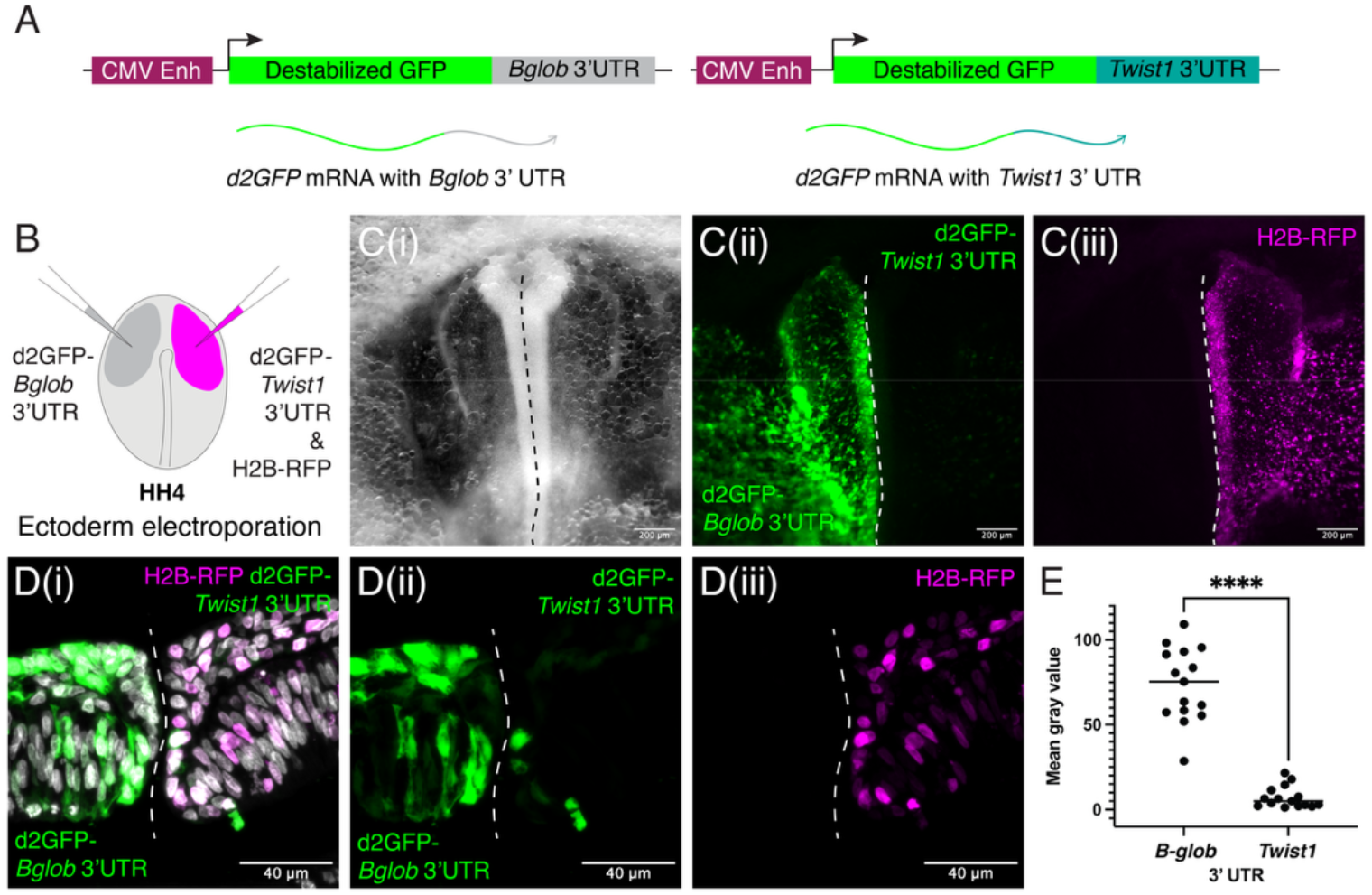
Post-transcriptional repression via the 3’ UTR of Twist1 occurs in the ectoderm. (A) Schematic of constructs used to test post-transcriptional repression. (B) Schematic representation of electroporation schema to test for post-transcriptional regulation of *Twist1* transcripts. The *beta-globin* 3’ UTR reporter was electroporated on the left side of the embryo, and the *Twist1* 3’ UTR reporter (with H2B-RFP plasmid as control) was electroporated on the right side of the embryo. (C) Single channel images of a representative embryo at HH8-9 showing (i) brightfield overview, (ii) GFP channel signal and (iii) RFP channel signal. The white dashed line denotes the midline of the embryo. N=15/15. (D) Representative section of cranial region at HH9 showing (i) composite image with DAPI, GFP and RFP signal, (ii) the GFP single channel image and (iii) the RFP single channel image. (E) Quantification of GFP signal intensity in segmented electroporated cells on either side of the embryo - measurements were taken from 3 independent embryos across 15 total sections. *Statistics calculated using paired t-test, p<0*.*0001*.

We observed bright GFP fluorescence in the neural tube and surface ectoderm on the left side of the HH8-10 embryo but little or no fluorescence in the same tissues on the right, *Twist1* 3’ UTR side of the embryo (*Figure 4C-E*). This result demonstrates that the *Twist1* 3’ UTR is strongly post-transcriptionally regulated in ectodermal tissues at this stage, effectively preventing *Twist1* transcript accumulation despite an active enhancer.

Our data reveal that *Twist1* expression is gated by a post-transcriptional mechanism in the neural tube, likely to prevent the premature mesenchymal differentiation of the CNCCs before reaching the pharyngeal arch environment. Therefore, we next sought to understand the precise mechanism by which the post-transcriptional repression is relieved to permit *Twist1* transcript abundance and acquisition of ectomesenchymal identity.

### A model for coordination of ectomesenchymal identity acquisition with spatial context

When the *Twist1* 3’ UTR reporter construct was electroporated throughout the epiblast and primitive streak of the embryo (*Figure 5A*), we observed GFP fluorescence preferentially in the ventral region of the developing head (*Figure 5B)* but little to no fluorescence dorsally. We also observed bright GFP fluorescence within the extraembryonic tissue (*Figure 5C*) as well as the somites (*Figure 5D*). Together these results suggest that the *Twist1* transcript is strongly destabilized in the neural tube and surface ectoderm, and that this destabilization is tissue specific.

**Figure 5:**
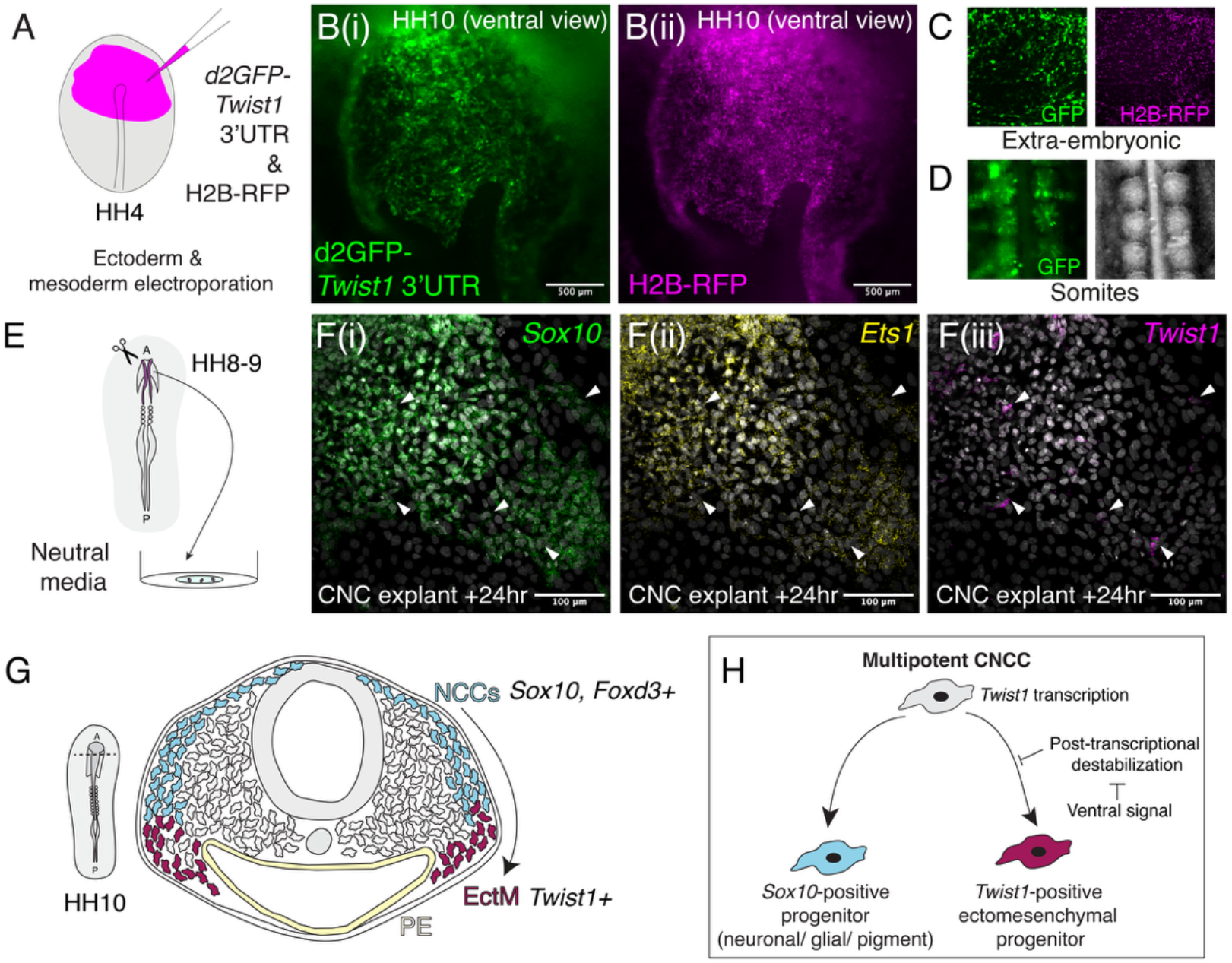
Post-transcriptional destabilization of Twist1 transcripts is context dependent. (A) Schematic representation of whole ectoderm electroporation. (B) Single channel images of a representative HH10 embryo in ventral view - (i) GFP signal and RFP signal. (C) Single channel images of extra-embryonic tissue showing GFP signal (left) and RFP signal (right). (D) Single channel images of trunk region showing GFP signal (left) in somite tissue – right image shows a brightfield overview. (E) Schematic of cranial neural crest explant – dorsal neural folds were dissected from HH8-9 embryos and explanted on fibronectin-coated glass dishes. (F) Multiplexed HCR staining for *Sox10, Ets1* and *Twist1* transcripts in CNC explants fixed after 24 hours of culture. Arrowheads indicate the few *Twist1* expressing cells in the explant. (G) Schematic summarizing model for emergence of ectomesenchymal identity in CNCCs. *Abbreviations - NCCs: neural crest cells, EctM: ectomesenchyme*.

Our observation of higher GFP fluorescence in ventral tissues suggests that *Twist1* transcript stability is higher within the ventral region of the embryo, consistent with the stabilization of *Twist1* transcripts in CNCCs late in their migration toward the pharyngeal arches. One possible explanation is that the extrinsic signaling environment of the ventral head influences transcript stability. To test whether extrinsic signals are required for *Twist1* transcript stabilization, we used an explant approach to assay neural crest gene expression when cultured in neutral media. Cranial neural folds were dissected from HH8-9 embryos, explanted on fibronectin in neutral media, and cultured for 24 hours (*Figure 5E*). In contrast to *in vivo* development, where these cells normally would upregulate *Twist1*, explants stained strongly for *Sox10* and *Ets1* but not *Twist1* (*Figure 5F*), suggesting that extrinsic cues from the embryonic environment are required for *Twist1* transcript stabilization.

Our observations support a model (*Figure 5G-H*) where *Twist1* transcription is activated in premigratory CNCCs within the neural tube, but *Twist1* transcripts remain highly unstable in the neural tube and dorsal head environment. As CNCCs migrate ventrally into the pharyngeal arches, they receive extrinsic signals that lead to the stabilization of *Twist1* transcripts, allowing rapid upregulation of *Twist1* and establishment of ectomesenchymal identity. Our model, decoupling enhancer priming and transcript stabilization, would ensure that ectomesenchymal commitment is acquired in the correct anatomical context. Such a mechanism may represent an evolutionary innovation that allowed vertebrates to precisely coordinate the spatial context of neural crest differentiation into ectomesenchyme. To test whether this regulatory logic is conserved across species, we next turned to interspecies UTR experiments.

### Interspecies 3’ UTR reporter experiments reveal conserved ectodermal post-transcriptional repression of Twist1 transcripts across vertebrates

To ask whether this phenomenon is species-specific, or if the 3’ UTRs from *Twist1* orthologues across vertebrates show the same destabilizing influence, we cloned additional constructs consisting of the d2GFP open reading frame followed by either the zebrafish (*Danio rerio*) *twist1a* 3’ UTR, the mouse (*Mus musculus*) *Twist1* 3’ UTR, or the frog (*Xenopus laevis*) *Twist1*.*L* 3’ UTR. All of these constructs produced a very similar result to the chicken construct upon electroporation into the chicken epiblast (*Figure 6A-D*), suggesting that the sequence elements responsible for destabilization of *Twist1* transcripts in the ectoderm are conserved. We also produced an equivalent construct containing the *twist1a-like* 3’ UTR from the non-vertebrate chordate *Ciona intestinalis*. Interestingly, this construct led to bright GFP fluorescence in the neural tube and surface ectoderm (*Figure 6E*), indicating that ectodermal repression of *Twist1* is absent outside of vertebrates and that sequence elements contributing to transcript destabilization evolved in vertebrates.

**Figure 6:**
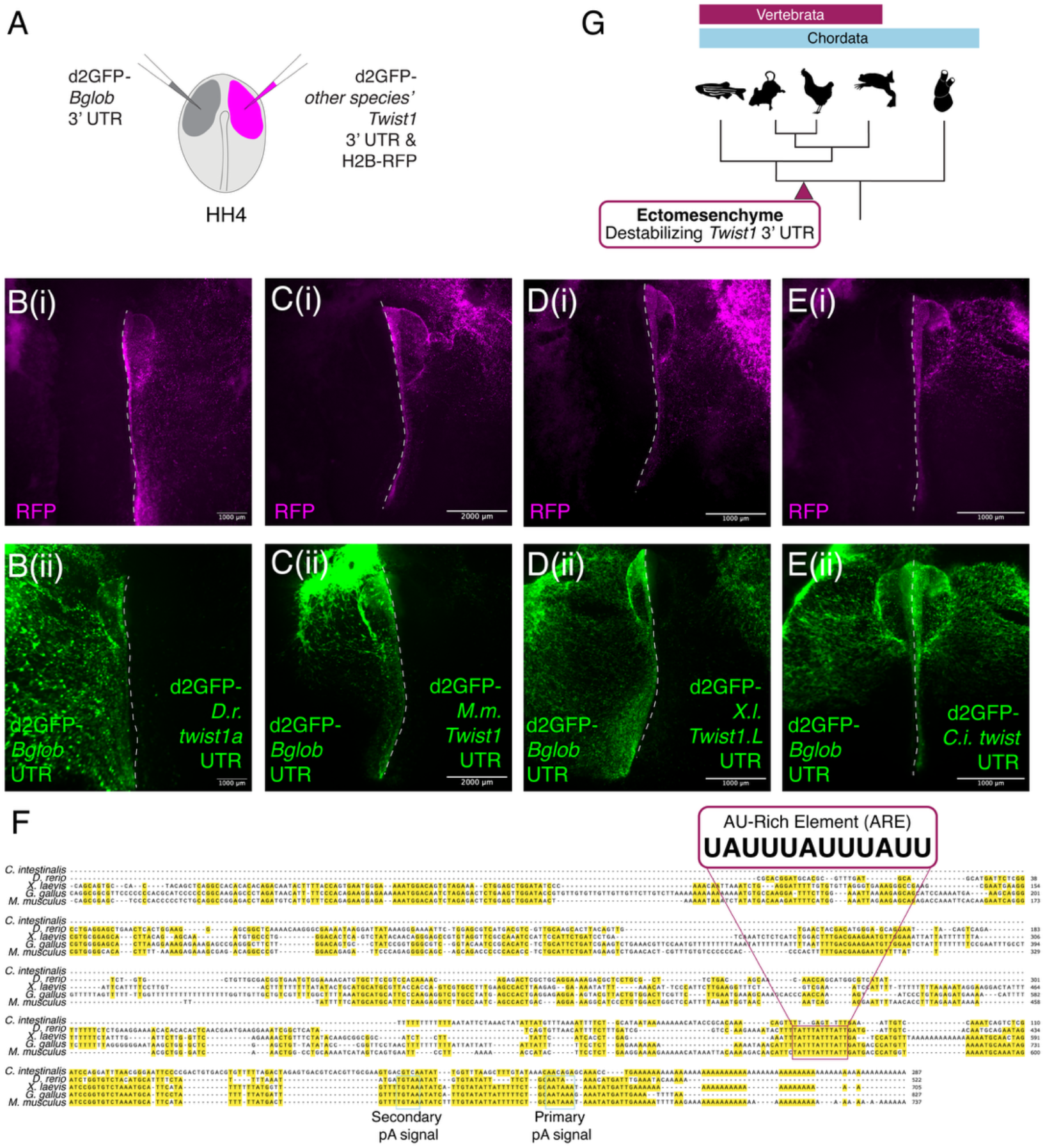
Post-transcriptional destabilization of Twist1 transcripts is vertebrate specific. (A) Schematic representing electroporation schema for interspecies UTR reporter experiments. The *Beta-globin* 3’ UTR reporter was electroporated on the left side of the embryo, and the *Twist1* 3’ UTR reporter from a different species was electroporated on the right side of the embryo with the H2B-RFP electroporation control. (B) *Danio rerio* 3’ UTR reporter, (C) *Mus musculus* 3’ UTR reporter, (D) *Xenopus laevis* 3’ UTR reporter and (E) *Ciona intestinalis twist1a-like* 3’ UTR reporter, each electroporated into the chicken embryo. For (B)-(E), panel (i) shows the H2B-RFP signal and panel (ii) shows the GFP signal. The dashed line in each image denotes the midline. Note expression of GFP on the right-hand side of the embryo that received the *Ciona* UTR construct but none of the embryos which received vertebrate 3’ UTR reporter constructs. (F) Multiple sequence alignment of *Twist1* 3’ UTR cDNA sequence between *C. intestinalis, D. rerio, X. laevis, G. gallus* and *M. musculus*. Yellow highlighted nucleotides are conserved between 3 or more sequences. The positions of the primary and secondary polyadenylation (pA) signals as well as a conserved AU-rich element (ARE) are indicated. Note that this ARE is conserved between all four vertebrate species but absent from *Ciona*. (G) Summary schematic – the sequence elements that destabilize the *Twist1* 3’ UTR evolved in vertebrates, facilitating the ability of the ectoderm to give rise to ectomesenchyme.

Given the high degree of functional conservation that we observed in our interspecies 3’ UTR reporter experiments, we next asked whether there are conserved sequence elements across vertebrate *Twist1* 3’ UTRs and absent from non-vertebrate *Twist* 3’ UTRs. We performed a multiple sequence alignment of *Twist1* 3’ UTR sequences from *C. intestinalis, D. rerio, X. laevis, G. gallus* and *M. musculus* (Figure 6F), finding that the most highly conserved sequence elements present in all four vertebrate species were concentrated toward the 3’ end of these UTR sequences. In particular, we noted the presence of an AU-Rich Element (ARE), <u>UAUUUAUUUAUU</u>, that is conserved across all four vertebrates but absent from the *C. intestinalis* 3’ UTR (*Figure 5F*). This ARE contains the nonamer, UUAUUUAUU, which has been previously shown to be the key sequence determinant mediating mRNA deadenylation and degradation (Lagnado et al., 1994; Zubiaga et al., 1995).

Together, these data demonstrate that post-transcriptional repression is an important contributor to proper *Twist1* expression and ectomesenchymal fate selection across vertebrates. More broadly, our results point to the evolution of the *Twist1* 3’ UTR as a molecular change that enabled ectoderm-derived neural crest cells to generate mesenchymal derivatives, an innovation that helped break classical germ layer boundaries, underpinning the emergence of the vertebrate head (Figure 6G).

## Discussion

### *Twist1* is an ectomesenchymal identity gene (not an EMT driver) in neural crest cells

*Twist* genes, which encode bHLH transcription factors, were first described in *Drosophila melanogaster* as key regulators of mesodermal identity and epithelial-to-mesenchymal transition (EMT) in ventral furrow cells (Leptin, 1991). Subsequently, the role of *Twist* genes in mesodermal identity was found to be widely conserved across metazoans including the vertebrate lineage (Chen and Behringer, 1995). Additionally, when *Twist1* was cloned in *Xenopus*, it was found to be expressed in CNCCs from neural plate border stages (stage 14.5+) and was subsequently implicated in the EMT of these cells (Hopwood et al., 1999; Lander et al., 2012). This led to the conception of *Twist1* as a key driver of EMT in neural crest cells, and consequently *Twist1* has been frequently included in Gene Regulatory Networks (GRNs) for neural crest cell specification at the neural plate border (Betancur et al., 2010b; Martik and Bronner, 2017; Simoes-Costa and Bronner, 2013).

Our data challenges this long-standing paradigm. In this study, we show that in avian and zebrafish embryos, *Twist1*/*twist1a* is not expressed in CNCCs until they are late in their migration, a timing that is inconsistent with a role for *Twist1* in driving neural crest EMT. Consistent with this, global and tissue-specific *Twist1* mouse mutants do not disrupt neural crest delamination, or migration into the frontonasal prominence and pharyngeal arches; instead, these CNCCs fail to acquire ectomesenchymal identity, maintaining inappropriate expression of *Sox10* in the pharyngeal arches and frontonasal prominence (Bildsoe et al., 2009; Soo et al., 2002). A very similar phenotype is observed in *twist1a/1b* double morphants or mutants in zebrafish (Das and Crump, 2012; Teng et al., 2018). Thus, *Twist1* is dispensable for EMT but essential for CNCC commitment to ectomesenchymal derivatives.

Interestingly, parallels exist in the context of *Drosophila* mesoderm development, where it has been shown that initiation of invagination and mesodermal identity emergence can be uncoupled. In the absence of *twist, snail* can drive invagination of prospective mesoderm cells, but these cells fail to upregulate mesodermal markers such as *tinman* (*Nkx2-5*) and *bagpipe* (*Bapx1*) (Ip et al., 1994). This result reinforces the idea that *Twist* genes in the mesoderm act primarily to reinforce cellular identity, consistent with our observations in neural crest cells.

Together, our results reframe our conception of *Twist1* in vertebrate neural crest biology. Rather than driving EMT, *Twist1* functions primarily as a gatekeeper for acquiring ectomesenchymal identity. Our findings not only resolve long-standing inconsistencies in the literature about the role of *Twist1* in early neural crest development but also highlight *Twist1* as a critical node in the regulatory circuitry that enabled ectoderm-derived neural crest cells to generate mesenchymal derivatives.

### Cell-intrinsic and extrinsic contributors to the ectomesenchymal cell fate decision

As discussed in the introduction, cranial neural crest cells express a distinct suite of transcription factors that have been shown to be sufficient to imbue trunk neural crest cells with the potency to form cartilage, an ectomesenchymal derivative (Simoes-Costa and Bronner, 2016). Additionally, extrinsic signaling cues from the head environment have been implicated in the emergence of ectomesenchymal identity and chondrocyte differentiation. For example, when quail CNCCs are dissociated at HH9 and cultured as isolated clones, they differentiate into glia, neurons, smooth muscle and melanocytes (Bhattacharya et al., 2018; Cohen and Konigsberg, 1975). Ectomesenchymal derivatives are underrepresented in these cultures, suggesting that signals from the embryonic environment may be important for their differentiation. One key signaling cue that is important for emergence of ectomesenchymal identity is pharyngeal endoderm-derived FGF ligands (Blentic et al., 2008), which recently have been shown to be sufficient for hESC-derived CNCCs to adopt ectomesenchymal identity and upregulate *Twist1* (Motoike et al., 2025).

Here, we provide mechanistic insight into how intrinsic and extrinsic contributions are integrated to drive ectomesenchymal fate acquisition. We propose that the accumulation of *Twist1* transcripts in cranial neural crest cells is driven by early enhancer activity in combination with potent transcript destabilization in the ectoderm. As CNCCs migrate ventrally within the developing head, extrinsic signals relieve this transcript destabilization, enabling rapid activation of the ectomesenchymal GRN program in the correct spatial context (*Figure 5G-H*). This mechanism reconciles previous observations including the absence of ectomesenchymal derivatives from cultures of CNCCs cultured *ex vivo*.

Consistent with previous studies, we identify an axially restricted *Twist1* enhancer that is active in cranial but not trunk neural crest cells; this restriction of enhancer activity may contribute toward the intrinsic difference in ectomesenchymal potency of cranial and trunk neural crest cells in amniotes. Together, our findings demonstrate that ectomesenchymal identity emerges through a two-tiered regulatory system: intrinsically through an axially restricted enhancer and *Twist1* transcription, and extrinsically through cues from the ventral head environment that stabilize *Twist1* transcripts to trigger mesenchymal differentiation.

### Post-transcriptional regulation in neural crest biology

Increasingly, post-transcriptional regulation has been implicated in neural crest biology (Bernardi et al., 2024; Bhattacharya et al., 2018; Copeland and Simoes-Costa, 2020; Eberhart et al., 2008; Hutchins et al., 2020; Keuls et al., 2023). For example, in mouse, the miRNA *miR-302* has been previously implicated in post-transcriptional regulation of *Sox9*, and prevention of precocious ectomesenchymal differentiation (Keuls et al., 2023). Another striking example of post-transcriptional regulation in the neural crest is the regulation of *Draxin* transcripts. *Draxin* encodes a Wnt antagonist that modulates cranial neural crest EMT and migration and is rapidly downregulated at the RNA level as NCCs delaminate (Hutchins et al, 2020). This rapid transcript downregulation is achieved post-transcriptionally, via degradation of transcripts upon delamination (Hutchins et al, 2020). Broadly, the importance of post-transcriptional regulation in neural crest gene regulatory networks is only beginning to be uncovered.

Our results demonstrate that *Twist1* transcripts in cranial neural crest cells are subject to potent post-transcriptional regulation in the avian early ectoderm, revealing a new layer of regulatory logic that gates the timing of ectomesenchymal identity acquisition. We note that *Twist1* has also been suggested to be under post-transcriptional regulation in mouse embryogenesis due to a non-concordance between the detection of *Twist1* transcript and TWIST1 protein (Gitelman, 1997). This raises the intriguing possibility that *Twist1* mRNA may also be regulated at the translational level either in addition to or in lieu of transcript degradation depending upon species context.

### Evolution of ectomesenchyme as a novel cell type

Ectomesenchyme is a vertebrate innovation; these cells defy germ layer theory, originating in the ectoderm and giving rise to mesenchymal (mesoderm-like) derivatives. We have shown that across multiple vertebrate lineages, the *Twist1* 3’ UTR sequence is sufficient to potently destabilize transcripts in the neural tube and surface ectoderm. Strikingly, the *twist1a-like* 3’ UTR sequence of the non-vertebrate chordate *Ciona intestinalis* lacks this destabilizing activity, underscoring that this regulatory property is a vertebrate-specific acquisition. We have also found that vertebrate *Twist1* 3’ UTRs contain an ARE, <u>UAUUUAUUUAUU.</u> This sequence element closely resembles AREs that have been shown to mediate deadenylation and subsequent degradation of mRNAs in other contexts; it is known to be recognized by RNA-binding proteins (e.g., Tristetraprolin and related proteins) that directly interact with components of the RNA deadenylation machinery (Lai et al., 2003; Tiedje et al., 2012).

Interestingly, a correlation has been described between 3’ UTR length and morphological complexity (number of cell types) in animal evolution (Chen et al., 2012). Consistent with this, we propose that the elaboration of the *Twist1* 3’ UTR sequence, through the introduction of destabilizing sequence motifs in vertebrates, was a critical step that facilitated the ability of the ectoderm to give rise to ectomesenchymal derivatives. By preventing precocious or inappropriate differentiation of these cells, this mechanism ensures fidelity of CNCC contributions to the head skeleton. Our findings highlight a novel regulatory innovation that – by restricting the emergence of ectomesenchyme to the correct anatomical context – may have facilitated the ability of neural crest cells to contribute to tissues associated with multiple germ layers.

## Author contributions

Experimental design: L.B., M.M., Experimental investigation: L.B., J.P., L.L., Data analysis: L.B., Computational Analysis: L.B, Manuscript – initial draft: L.B., Manuscript – editing: L.B., M.M., L.L., J.P.

## Acknowledgements

The authors thank Erica Hutchins for gifting the destabilized GFP (d2GFP) β-globin 3’ UTR construct. We thank the animal care team in the UC Berkeley fish facility, including Tyler Mentley, Frances Campbell, and Lindsey Arenson. The authors would also like to thank members of the Martik lab for helpful discussions and Hannah Van Mullem and Rekha Dhillon-Richardson for critical reading of the manuscript.

## Funding

This work was supported by an EMBO postdoctoral fellowship to L.C.B. and NIH K99/R00HD100587, NIH DP2HL173858, and UC Berkeley start-up funds awarded to M.L.M.

## Competing interests

The authors declare no competing financial interests.

## Materials and methods

### Animal and embryo husbandry

Fertilized chicken (Gallus gallus) eggs were sourced from a local farm (Petaluma Egg Farm, CA) and incubated in a humidified incubator at 37°C to obtain embryos of the desired stage according to Hamburger and Hamilton (1951). Embryos were fixed in 4% PFA in PBS overnight at 4°C and washed in PBST.

Wild-type AB strain adult zebrafish (Danio rerio) were maintained in a 14/10 h light/dark cycle at 28 °C and fed twice daily. All experiments were approved and conducted in compliance with UC Berkeley Institutional Animal Care and Use Committee (IACUC) protocol AUP-2021-03-14107-1. Embryos were collected and maintained at 28.5°C in egg water (Westerfield, 2007), and staged according to somite number (Kimmel et al., 1995). Following staging, zebrafish embryos were dechorionated with 0.5 mg/mL pronase in egg water for approximately 10-15 minutes, rinsed 3x with egg water to remove residual pronase, and fixed in 4% PFA in PBS for 1-2 hours at room temperature or overnight at 4°C.

### Electroporation

For *ex ovo* electroporation, embryos were cultured on filter paper according to established protocols (Chapman et al., 2001). Agar-albumen plates were prepared as follows: 0.6g of agarose was dissolved in 100 ml of saline solution, cooled to 37°C and then combined with 100 ml of thin albumen freshly obtained from eggs as well as Penicillin-Streptomycin solution (ThermoFisher product 15140122) to 1X concentration. Plasmid solution (each plasmid at 2µg/µl in Ringer’s solution with Brilliant Blue dye) was microinjected beneath the epiblast of the HH4 embryo and a 5.6V current applied across the dorsoventral axis of the embryo (5 pulses of 50ms each). For in ovo electroporation, plasmid solution (each plasmid at 2µg/µl in Ringer’s solution with Brilliant Blue dye) was microinjected into the lumen of the neural tube. A small volume of Ringer’s solution was dropped onto the embryo, and an electrical current was passed from the left to the right of the embryo (21V current, 3 pulses of 50ms each). Eggs were sealed with tape and allowed to incubate for an additional 16-20 hours.

### Neural crest explants

Mattek glass bottomed imaging dishes were coated with human fibronectin protein (Biotechne, 1918-FN-02M) as follows: a 25µg/ml fibronectin solution was made up in PBS using a 1:40 dilution of 1mg/mL stock solution. 100µl of fibronectin solution was pipetted onto the glass surface, then the dish was incubated at 37°C for 1-3 hours. Fibronectin solution was removed, and the dish was allowed to fully dry at RT. Dorsal neural folds were microdissected in PBS from chicken embryos at HH8-9, then placed onto fibronectin coated glass dishes containing GMEM (Gibco) + 10% Fetal Bovine Serum (FBS) + 1% Penicillin-Streptomycin solution. Explants were incubated at 37°C with 5% CO_2_ for 24 hours, then fixed in 4% PFA at RT for 20 minutes.

### Enhancer construct cloning

Putative enhancer sequences were ordered as g-blocks with appropriate overhangs (Twist Bioscience) and cloned into the published Sox10 E2>GFP vector (Betancur et al., 2010a). The Sox10 E2 enhancer sequence was removed by successive restriction digestion reactions with KpnI-HF (NEB) and BglII (NEB), then g-blocks and digested plasmid were assembled by Gibson assembly. For scrambled enhancer construct analyses, the predicted transcription factor binding site within the *Twist1*+225kb enhancer sequence was scrambled using a random DNA sequence scrambler (Shuffle DNA tool). To validate that the scrambling of this sequence did not remove or create any additional transcriptional factor binding sites other than the target, the scrambled enhancer sequence was then scanned for TF binding motifs and the output compared to the unscrambled enhancer output. All putative and validated enhancer sequences are included in *Supplementary Table 2*.

### 3’ UTR construct cloning

The destabilized GFP (d2GFP) β-globin 3’ UTR construct was a gift from Erica Hutchins (Hutchins et al., 2021). To replace the β-globin 3’ UTR with the *Gallus gallus Twist1* 3’ UTR, the *G. gallus Twist1* 3’ UTR sequence (827bp, *Supplementary Table 3*) was ordered as a g-block with appropriate overhangs to assemble into the plasmid. The *β-globin* 3’ UTR plasmid was digested with NotI and BsaBI, then Gibson assembled with the g-block. The same process was used to clone the *Danio rerio twist1a, Mus musculus Twist1, Xenopus laevis Twist1*.*L* and *Ciona intestinalis twist* 3’ UTR reporter constructs.

### Hybridization chain reaction (HCR v3.0)

Chicken embryos were fixed in 4% paraformaldehyde in PBST for two hours at room temperature or overnight at 4°C, then dehydrated through a methanol/PBST series to 100% methanol and stored at - 20°C. Embryos were rehydrated through a methanol/PBST series and then washed several times in PBST at room temperature. Chicken embryos were digested with proteinase K (NEB, 1mg/ml in PBST) for 1 minute per Hamburger Hamilton stage, quick-washed in PBST and then postfixed in 4% PFA for 20 minutes. Embryos were then washed 3 times in PBST for 10 minutes, twice in 5X SSCT and then pre-hybridized in HCR V3.0 hybridization buffer for 30 minutes at 37°C. Probes were applied at 0.8pmol per 500µl hybridization buffer and embryos were incubated overnight at 37°C. The next day, embryos were washed 4 times for 15 minutes each with HCR wash buffer at 37°C. Embryos were then washed three times with 5X SSCT at room temperature, and then pre-incubated in HCR amplification buffer for 5-30 minutes at room temperature. HCR hairpins were snap-cooled by heating to 95°C for 90 seconds, then cooled to room temperature in darkness. Hairpins were added to HCR amplification buffer at 4µl per 500µl of buffer. Embryos were then incubated in hairpin solution overnight in darkness at room temperature. Finally, embryos were washed 5-8 times (10 minutes each) in 5X SSCT at room temperature and DAPI stained. For chicken explants, the above protocol was followed with the following modifications: fixation was for 20 minutes at RT in 4% PFA, dehydration was in ethanol rather than methanol, and proteinase K treatment and post-fixation were omitted. For zebrafish embryo HCRs, the above protocol was followed with the following modifications: dehydration was in ethanol rather than methanol, and proteinase K treatment and post-fixation were omitted. Probe concentrations were 2 pmol of probe in 500µL of probe hybridization buffer. Hairpin concentrations were 6 pmol per hairpin (2µL each) per 500µL of amplification buffer. All probes, hairpins, and buffers were purchased through Molecular Instruments.

### Cryosectioning

Pre-stained chicken embryos were washed in PBS, equilibrated in 5% sucrose in PBS for 30 minutes, and then in 15% sucrose in PBS overnight at 4°C. Gelatin solution (7.5% gelatin, 15% sucrose in 0.5X PBS) was melted at 37°C and applied to embryos, which were then incubated at 37°C for 3-4 hours to allow equilibration. After this incubation, individual embryos were mounted in coffin molds in gelatin solution which was allowed to solidify with embryos in the appropriate orientation. After fully solidifying, blocks were flash-frozen in liquid nitrogen, stored at -80°C and then cryosectioned at 12µm on a Leica CM3050 cryostat. Slides were incubated in PBS in a coplin jar in a 37°C water bath to dissolve gelatin, then coverslipped with fluoromount (Southern Biotech) and allowed to set overnight at room temperature before confocal imaging.

### Plastic sectioning of zebrafish embryos

Following whole-mount HCR, zebrafish embryos were washed 3 times for 5 minutes in PBST and manually deyolked by puncturing the yolk with forceps and gently pipetting up and down to remove yolk granules. Embryos were then embedded in plastic using the JB-4 embedding kit and following a previously published protocol (Sullivan-Brown et al., 2010). Embedded embryos were sectioned on a Leitz 1512 microtome at a thickness of 7-10µm, and sections were adhered to slides by being placed on 20-40µL droplets of boiled ddH2O on a 50°C slide warmer. Once the water fully evaporated, slides were mounted in fluoromount (Invitrogen), coverslipped and left to set overnight before confocal imaging.

### Confocal imaging

For whole-mount zebrafish HCRs, embryos were mounted in 0.42μm canyon molds and imaged with a 20X dipping lens on a Zeiss 780 LSM upright confocal microscope. For sectioned HCRs, images were taken using a 40X objective lens. Image analysis was performed using Fiji, and all images are displayed as summed intensity projections.

### Computational analysis

#### Hi-ChIP

Raw sequencing reads from previously published H3K27ac Hi-ChIP datasets from HH9 dissected neural folds and whole embryos (Azambuja & Simoes-Costa, 2021) were downloaded from the European Nucleotide Archive (accession numbers SRR11719979-SRR11719980).

Sequencing reads were mapped to the galGal6 genome assembly using HiC-Pro (Servant et al., 2015), then peaks were called and loop anchoring performed using Hichipper (Lareau & Aryee, 2018a). Output bedpe files from Hichipper were then read into R and Diffloop (Lareau & Aryee, 2018b) was used to filter contacts (minimum loop size 1.5kb, minimum of 5 replicates in 3 samples) and perform statistical analysis on neural crest enriched and depleted loops relative to whole embryo samples. To visualize enhancer-promoter contacts, the Sushi package (Phanstiel et al., 2014) was utilized in R.

#### Cut & Run

Raw sequencing reads from previously published TFAP2A (Rothstein and Simoes-Costa, 2020), TFAP2B (Rothstein and Simoes-Costa, 2020) and H3K27ac (Bhattacharya et al., 2020) Cut & Run experiments were downloaded from the European Nucleotide Archive (accession numbers SRX5402880-SRX5402881; SRX5402884-SRX5402885; SRX7388167-SRX7388168). Raw reads were mapped to the galGal6 genome assembly using Bowtie2 (Langmead & Salzberg, 2012).

#### ATAC-Sequencing

Raw sequencing reads from previously published HH10 and HH12 sorted CNC ATAC-sequencing experiments (Hovland et al., 2022) were downloaded from the European Nucleotide Archive (accession numbers SRR13295154-SRR13295157). Adaptor sequences (forward: CTGTCTCTTATACACATCT, reverse: AGATGTGTATAAGAGACAG) were trimmed from reads using cutadapt (Martin, 2011). Trimmed reads were then mapped to the galGal6 reference genome using Bowtie2 (Langmead & Salzberg, 2012). Replicates for each sample were combined using samtools merge function (Li et al., 2009), then bigwig files were generated for visualization purposes using the bamCoverage function from deepTools (Ramirez et al., 2016).

#### Visualization of multimodal sequencing datasets

To visualize ATAC and Cut & Run datasets together over a chromosomal interval of interest, the package PlotGardener in R was utilized (Kramer et al., 2022).

#### Sequence conservation

The Evolutionarily Conserved Region (ECR) Browser (Ovcharenko et al., 2004) was used to survey the *Twist1* genomic locus for non-coding sequences that are evolutionarily conserved among vertebrate genomes. Candidate putative enhancer sequences were converted from the galGal3 genome assembly coordinates to galGal6 coordinates using the UCSD Genome Browser.

#### Transcription factor binding site analysis

Candidate and/or validated enhancer sequences were scanned for transcription factor binding motifs using the Find Individual Motif Occurrences (FIMO) tool from MEME Suite (Grant et al., 2011). Sequences were analyzed against a library of motifs (all redundant vertebrate motifs) downloaded from the JASPAR database (Rauluseviciute et al., 2024).

#### Image quantification

To quantify GFP signal intensity from d2GFP 3’ UTR bilateral electroporations, sectioned embryos were imaged on a confocal microscope using a 40X oil immersion objective under constant laser intensity and gain settings. Images were processed using FIJI (Schindelin et al., 2012) as follows: each Z-stack was converted to a summed slices intensity projection, then the ‘Threshold Color’ tool was used to segment RFP-positive pixels (representing successfully electroporated cells on the experimental side of the embryo). The ‘Measure’ tool was used to measure mean pixel intensity in the GFP channel within the segmented RFP-positive region. Next, the image was cropped to only include the control *Beta-globin* 3’UTR side of the embryo, then the ‘Threshold Color’ tool was used to segment and select all GFP-positive pixels (representing successfully electroporated cells). The ‘Measure’ tool was used to measure mean pixel intensity within the selected region. Finally, the measurements of mean pixel intensities on the left and right halves of the embryo were collated across multiple replicates and GraphPad Prism was used to plot data and perform a paired t-test.

To quantify RFP and GFP signal intensity from widefield images of bilateral scrambled enhancer electroporations, three measurements of corrected total cell fluorescence were taken from the neural tube (NT) and migratory neural crest (NC) regions on either side of the embryo (left and right). These measurements were normalized for each embryo (image) by dividing by the highest value; with this normalization performed separately for measurements of the neural tube and the migratory neural crest regions, then plotting and unpaired t-tests were performed in GraphPad Prism.

---

**Figure S1:**
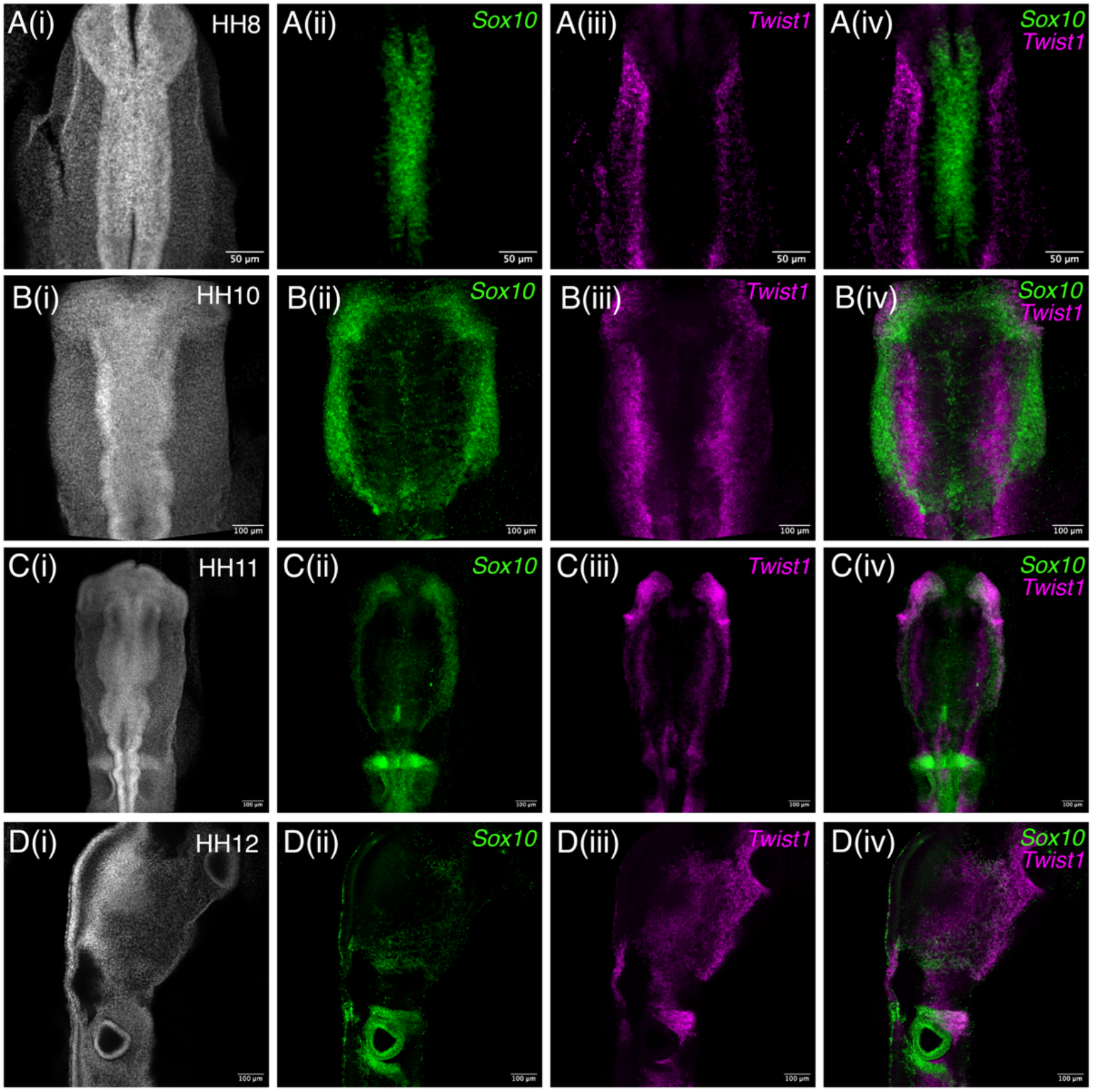
Wholemount HCR staining for Sox10 and Twist1 in chicken. Multiplexed *in situ* hybridization (V3.0 HCR) for *Sox10* and *Twist1* transcripts in wholemount embryos at (A) HH8, (B) HH10, (C) HH11, and (D) HH12. All images shown are summed slice intensity projections, with (i) of each panel showing DAPI nuclear staining, (ii) showing *Sox10* single channel image, (iii) showing the *Twist1* single channel image and (iv) showing the composite *Sox10* and *Twist1* image. *Scale bars in (A) represent 50μm and in (B)-(D) represent 100μm*.

**Figure S2:**
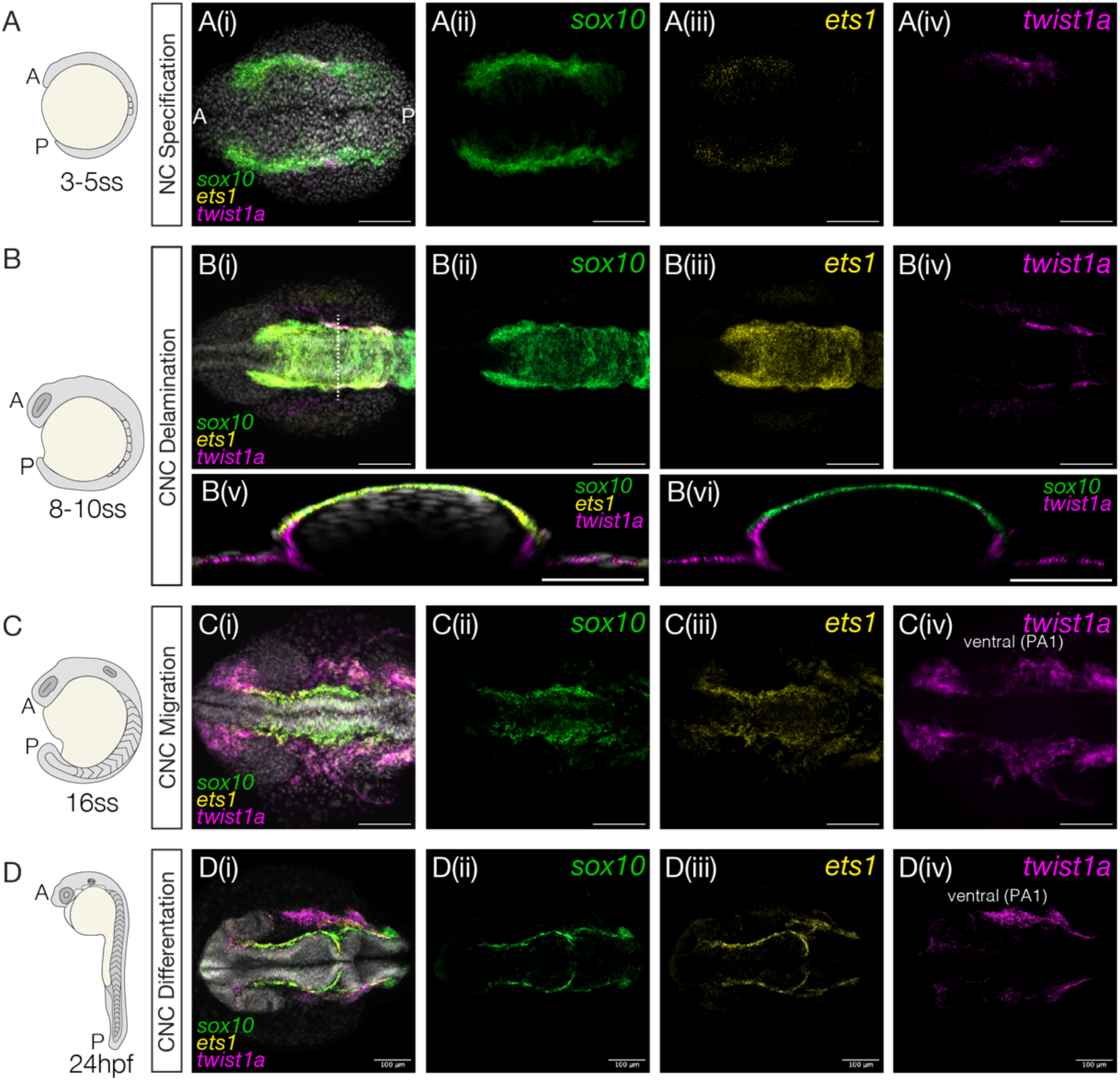
Wholemount HCR staining for sox10, ets1 & twist1a in zebrafish. (A)-(D) Schematic of zebrafish embryos during critical neural crest developmental stages (specification, delamination, migration, and differentiation) followed by summed slice intensity projections of multiplexed HCR staining for *sox10, ets1*, and *twist1a*. (i) shows a composite image with DAPI, *sox10, ets1* and *twist1a*, and (ii)-(iv) are single-channel images for *sox10, ets1*, and *twist1a*, respectively. (v) - (vi) is a resliced image of the same embryo in (B) along the yz-plane to show the distribution of *sox10* and *twist1a* transcripts. (v) shows a composite image with DAPI, *sox10, ets1* and *twist1a*, and (vi) is a double-channel image of *sox10* and *twist1a*. The axial level of the reslice is indicated by the dotted line drawn on (Bi). *Acronyms: A, anterior; P, posterior; PA1, pharyngeal arch 1. Scale bars are 50μm for resliced images (Bv-vi) and 100μm for all other images*.

**Figure S3:**
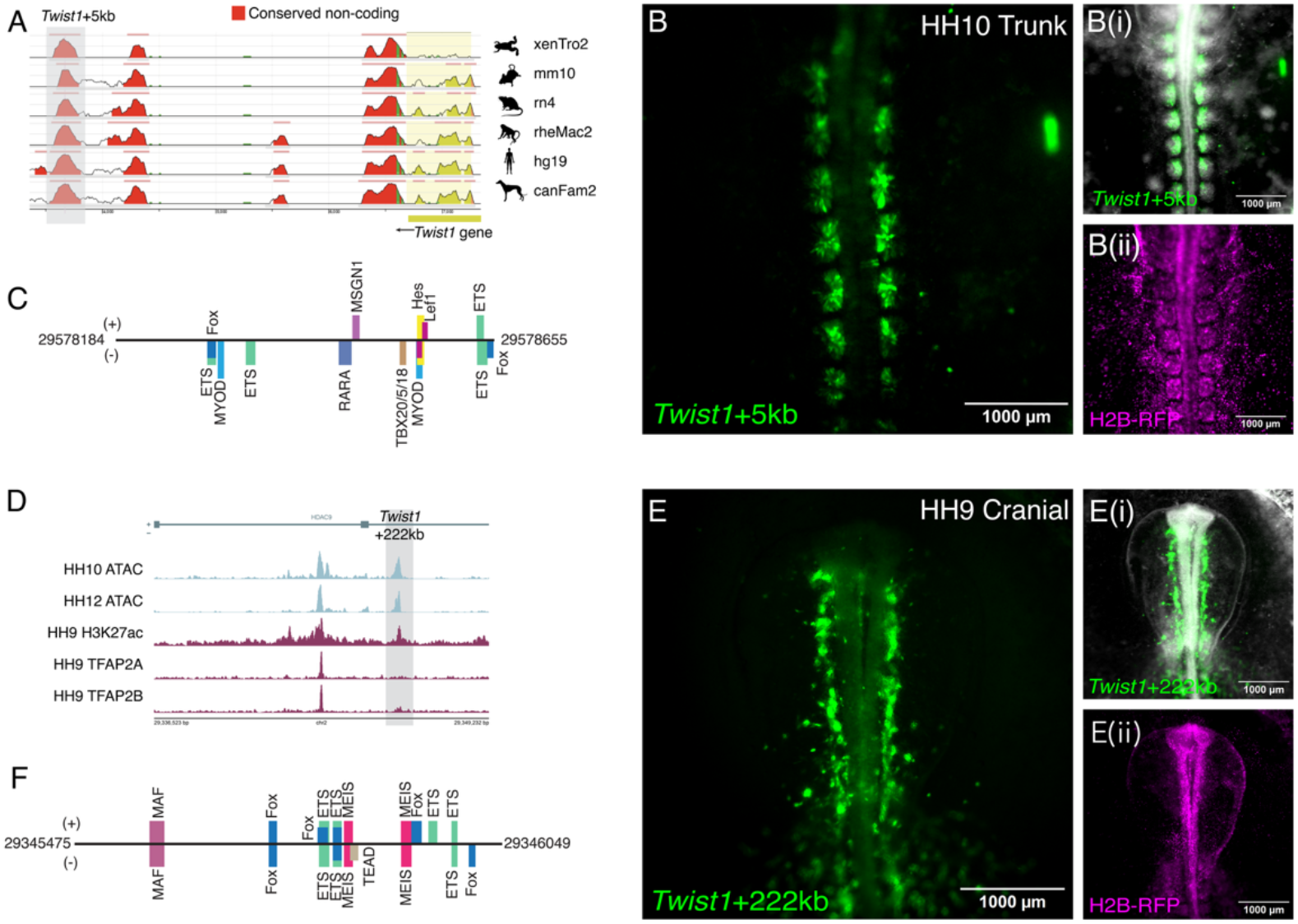
Validated mesodermal enhancers for Twist1. (A) Conserved sequence plot from the ECR Browser in the genomic region just downstream of the *Twist1* gene (base genome galGal3), with red peaks indicating regions of sequence homology between vertebrate genomes and galGal3 genome sequence. The *Twist1*+5kb enhancer peak is indicated by the grey shaded box. (B) Fluorescence images showing (B) GFP fluorescence, (B(i)) a composite brightfield and GFP fluorescence image and (B(ii)) H2B-RFP fluorescence in a representative embryo at HH10. GFP fluorescence is detected in the medial portion of the somites. (C) Box diagram of the *Twist1*+5kb enhancer with predicted transcription factor binding sites indicated. (D) Genome tracks showing chromatin accessibility, H3K27ac and TFAP2 transcription factor binding in the region encompassing the *Twist1*+222kb enhancer. *Note that this enhancer is contained within the same Hi-ChIP contact with the Twist1 promoter region as Twist1+225kb* (see main text *Figure 2*A-B). (E) Fluorescence images showing (E) GFP fluorescence, (E(i)) a composite brightfield and GFP fluorescence image and (E(ii)) H2B-RFP fluorescence in a representative embryo at HH9. GFP fluorescence is detected in cranial mesoderm close to the midline. (F) Box diagram of the *Twist1*+222kb enhancer with predicted transcription factor binding sites indicated.

## Notes

### Competing Interest Statement

The authors have declared no competing interest.

